# A general deep hybrid model for bioreactor systems: combining first Principles equations with deep neural networks

**DOI:** 10.1101/2022.06.07.495118

**Authors:** José Pinto, Mykaella Mestre, Rafael S. Costa, Gerald Striedner, Rui Oliveira

## Abstract

Numerous studies have reported the use of hybrid semiparametric systems that combine shallow neural networks with mechanistic models for bioprocess modeling. Here we revisit the general bioreactor hybrid modeling problem and introduce some of the most recent deep learning techniques. The single layer networks were extended to multi-layer networks with varying depths and combined with First Principles equations in the form of deep hybrid models. Deep learning techniques, namely the adaptive moment estimation method (ADAM), stochastic regularization and depth-dependent weights initialization were evaluated. Modified sensitivity equations are proposed for the computation of gradients in order to reduce CPU time for the training of deep hybrid models. The methods are illustrated with applications to a synthetic dataset and a pilot 50 L MUT+ *Pichia pastoris* process expressing a single chain antibody fragment. All in all, the results point to a systematic generalization improvement of deep hybrid models over its shallow counterpart. Moreover, the CPU cost to train the deep hybrid models is shown to be lower than for the shallow counterpart. In the pilot 50L MUT+ *Pichia* pastoris data set, the prediction accuracy was increased by 18.4% and the CPU decreased by 43.4%.

**Highlights:**

- Shallow hybrid models have been widely used for bioprocess modeling and optimization
- Non-deep training using e.g. the Levenberg – Marquardt method, cross-validation and indirect sensitivity equations have been the methods of choice
- Deep learning with ADAM, stochastic regularization and indirect sensitivity significantly reduces the training CPU
- The generalization capacity of deep hybrid models systematically outperforms that of shallow hybrid models

## INTRODUCTION

The first steps towards the integration of mechanistic abstraction and neural networks in process systems engineering were taken in the early 90’s with the pioneering works of (Psichogios and Ungar, 1992; Su and Mcavoy, 1993; Schubert et al., 1994) and (Thompson and Kramer, 1994). The main motivation was to overcome neural networks limitations, namely the i) inability to comply with process constraints, ii) the tendency for data overfitting, and iii) the poor predictive power outside the training-validation domain. Thompson and Kramer (1994) framed this problem as hybrid semi-parametric systems, whereby parametric functions with fixed structure stemming from prior process knowledge (e.g., macroscopic material balance equations) are combined in series or in parallel with nonparametric functions (e.g. neural networks) identified from process data. Numerous bioprocess hybrid modeling studies followed (e.g. (Preusting et al., 1996; van Can et al., 1998; Chen et al., 2000; Galvanauskas et al., 2004; Oliveira, 2004; Teixeira et al., 2007; Fiedler and Schuppert, 2008; von Stosch et al., 2011; Ferreira et al., 2014; Pinto et al., 2019; O’Brien et al., 2021; Bayer et al., 2021) highlighting the advantages of the technique, which may be summarized as a more rational usage of prior knowledge eventually translating into more accurate, transparent and robust process models.

The vast majority of hybrid modeling studies explored the combination of conservation laws and shallow neural networks (see review by (von Stosch et al., 2014)). Recent advances in deep learning have however demonstrated that neural networks with multiple hidden layers (deep networks) are advantageous over their shallow counterparts. Shallow and deep networks are both universal function approximators, but deep networks are able to approximate compositional functions with exponentially lower number of parameters and sample complexity (Delalleau and Bengio, 2011; Eldan and Shamir, 2016; Liang and Srikant, 2017) and are less prone to overfitting (Mhaskar and Poggio, 2016). The shift from shallow to deep network architectures has been triggered by the development of stochastic gradient descent training algorithms, particularly the ADAM method (Kingma, 2014). ADAM is a first-order gradient-based method for stochastic objective functions based on adaptive estimates of lower-order moments. The data subsampling along with the learning rate adaptation at each iteration resulted in a simple and robust training method that is less sensitive to gradient attenuation and to the convergence to local optima. Stochastic regularization based on weights dropout has been shown to effectively avoid overfitting in deep learning (Hinton et al., 2012; Srivastava et al., 2014). Stochastic regularization is frequently associated with stochastic gradient descent methods to prevent overfitting and to improve generalization properties (Koutsoukas et al., 2017).

Only very recently the deep learning advances are penetrating the hybrid modeling field. Bangi and Kwon (2020) proposed a hybrid model for a hydraulic fracturing process that combines a First Principles model with a deep neural network. A fully connected network with 5 layers (1×20×20×20×1), hyperbolic tangent activation (*tanh*) in the 3 hidden layers and linear activation in the input/output layers, was adopted. The Levenberg–Marquardt algorithm and finite difference-based sensitivity analysis were adopted to train the hybrid model. The resulting hybrid model had superior extrapolation properties compared to a purely data-driven deep neural network model.

In this study, we revisit the general bioreactor hybrid model (Oliveira, 2004; Teixeira et al., 2007; von Stosch et al., 2011; Ferreira et al., 2014; Pinto et al., 2019) and extend it to deep learning. More specifically, we explore the adoption of deep learning techniques in a hybrid semiparametric modeling context, such as deep feedforward neural networks with varying depths, the rectified linear unit (ReLU) activation function, dropout regularization of network weights, and stochastic training with the ADAM method. These techniques are applied to two case studies and are benchmarked against the traditional shallow hybrid modeling approach.

## MATERIALS AND METHODS

### General deep hybrid model for bioreactor systems

A stirred tank bioreactor can be generically represented by the hybrid model structure of Figure 1. The dynamics of process state variables are modelled by a system of ordinary differential equations (ODEs) derived from macroscopic material balances and/or intracellular material balances and/or other physical assumptions. These equations take the following general form:

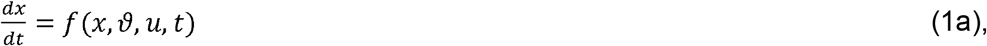

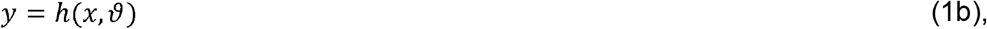

with *t* the independent variable time, *x*(*t*) the process state vector, *u*(*t*) the vector of external inputs (feed rates, temperature, pH, ionic strength, osmolality, etc), *ϑ* a vector of process variables with unknown defining functions, and *y* the vector of measured variables. Eq (1a,b) are the state-space model and measurement model respectively. The functions *f*(.) and *h*(.) are of parametric nature thus with fixed structure stemming from prior knowledge. Some relevant bioprocess variables may be less defined in terms of explanatory mechanisms and/or rely on loose assumptions. Typical examples are biological reaction kinetics or product quality attributes, which are difficult to establish on a mechanistic basis. In the general hybrid model, such properties are defined as loose functions, *ϑ*(·) (typically of the process state and external inputs), with unknown structure, i.e. nonparametric functions without physical meaning. Among the many possibilities to define *ϑ*(·), the preferred approach in the hybrid modeling literature has been by far the feedforward perceptron networks with 3 layers only (see review by (von Stosch et al., 2014)). In the present study, the more general case of deep multilayer perceptron networks with arbitrary number of *nh* hidden layers is explored, stated as follows:

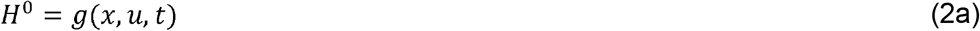

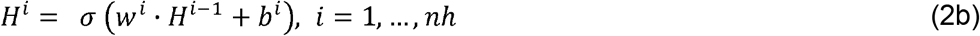

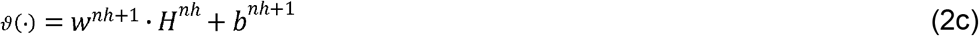

The input layer (Eq. 2a) typically receives information of the state variables, *x*, and/or external inputs, *u* (temperature, pH, etc,…) and/or process time, *t*. An optional non-linear pre-processing function, *g*(*x, u, t*), may sometimes facilitate the identification of *ϑ*(·), as for example concentration ratios are set as inputs to the neural network or other normalization rules (see (von Stosch et al., 2016; Gnoth et al., 2008; Gnoth et al., 2010)). Then follows *nh* hidden layers (Eq. 2b) with *σ*(·) the nodes transfer function (in this study either the *tanh* or the *ReLU*). Finally, the output layer has linear nodes (Eq 2c). The parameters *w*= {*w*^1^,*w*^2^,*…, w*^*nh*+1^} and *b*= {*b*^1^, *b*^2^,*…, b*^*nh*+1^} are the nodes connection weights that need to be identified from data during the training process. Presuming that initial conditions *x(t) = x*_0_ are given and that the network weights *ω* = {*w,b*} are defined, the deep hybrid model can be solved by numerical integration as an Initial Value Problem. In the present study, a Runge-Kutta 4^th^ order ODE solver was adopted to integrate the system (Eqs. 1-2) and compute *x, y* and *ϑ* over time. All the code was implemented in MATLAB on a computer with Intel(R) Core(TM) i7-9750H CPU @ 2.60 GHz 2.59 GHz, and 16 GB of RAM.

**Figure 1.**
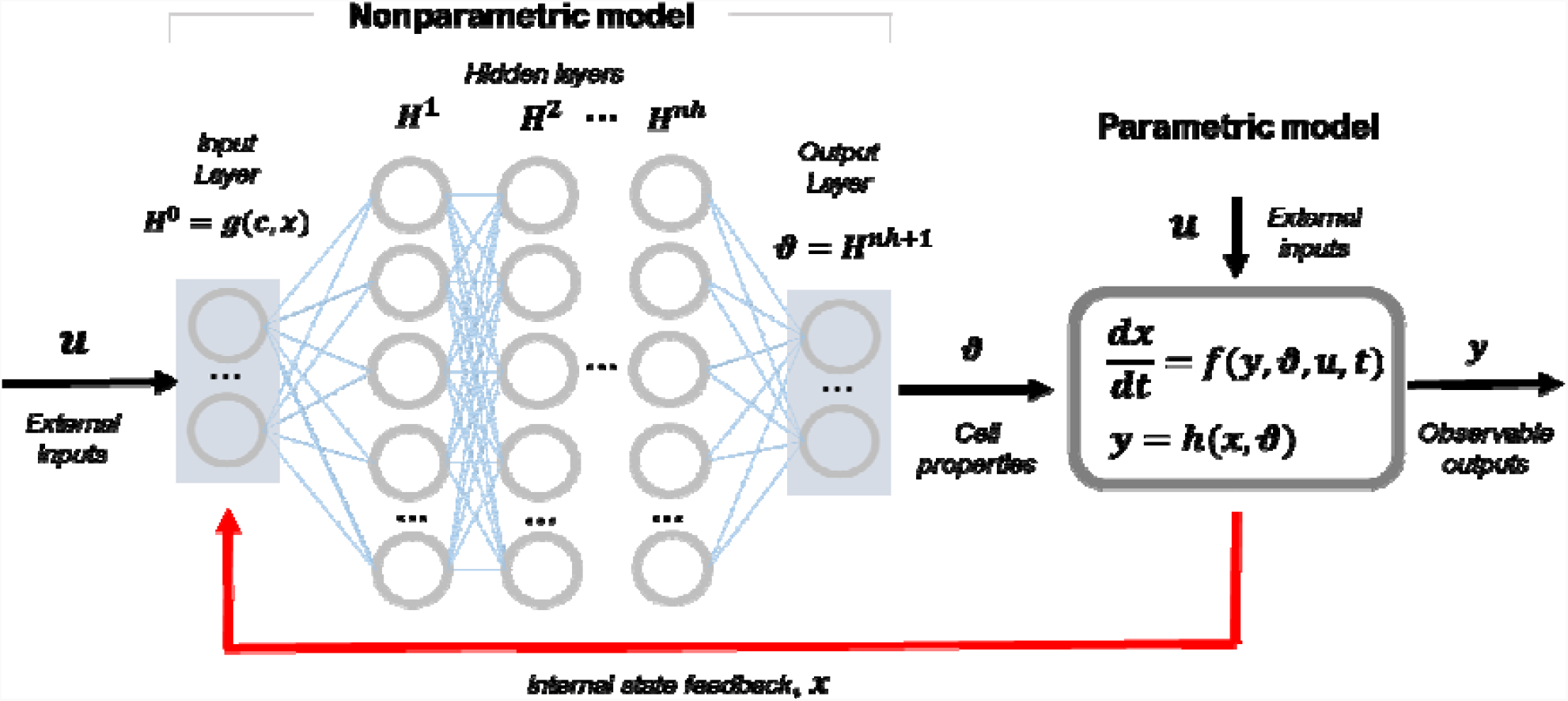
Schematic representation of the general deep hybrid model for bioreactor systems. The model is dynamic in nature with state vector, **x**, and observable outputs **y**. The model has a parametric component (functions f(.) and h(.)) with fixed mathematical structure determined by first Principles (typically material/energy balance equations). Some cellular properties are modelled by a deep feedforward neural network with multiple hidden layers as function of the process state, **x**, and external inputs, **u** (nonparametric model component with loose structure that must be identified from process data given the absence of explanatory mechanisms for that particular part of the model)

### Training method

#### Standard non-deep method

Hybrid bioreactor models are typically trained by indirect supervised learning with cross-validation to avoid overfitting (e.g., (Psichogios and Ungar, 1992; Oliveira, 2004; Pinto et al., 2019; von Stosch et al., 2014)). The data are partitioned in a training subset (for parameter estimation), a validation subset (stop criterion to avoid overfitting) and a test subset (to assess the predictive power). Partitioning is typically performed batch wise with the amount of data allocated in each partition depending on the objective of the study and on the amount of data available. The optimization of network parameters is performed over the training set only in a weighted least-squares sense:

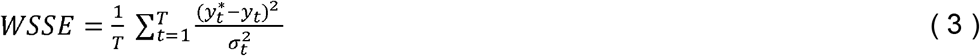

with *T* the number of training patterns, 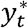the measured variables at time *t, y*_*t*_ the corresponding model prediction and *σ*_*t*_ the measurement standard deviation. This method is called indirect because the loss function is not directly linked to the neural network outputs, *ϑ*. The Levenberg-Marquardt method (LMM), has been shown to be very effective to solve the indirect training problem (Eqs. 1-3) in the case of shallow hybrid models (Schubert et al., 1994; Oliveira, 2004). The LMM has also been used in a recent deep hybrid modeling study (Bangi and Kwon, 2020). The LMM method convergence is improved if the sensitivity equations are applied to calculate the loss function gradients instead of numerical gradients (e.g. (Psichogios and Ungar, 1992; Schubert et al., 1994; Oliveira, 2004)). The sensitivity equations for the general hybrid model have the following structure (for simplicity it is assumed that *y= x*):

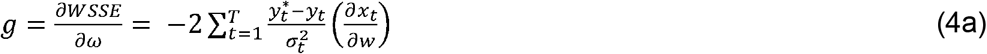

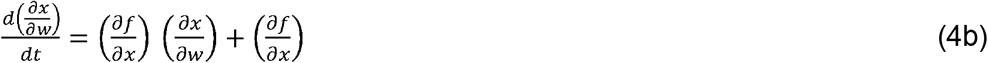

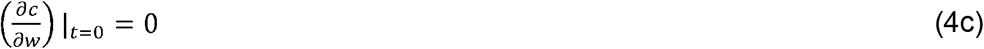

For more details regarding the sensitivity equations in a hybrid modeling context see (Psichogios and Ungar, 1992; Oliveira, 2004). The integration of the sensitivity equations was performed in this study with a Runge-Kutta 4^th^ order ODEs solver.

#### Stochastic adaptive moment estimation (ADAM) with semi-direct sensitivities

An important goal of this study is to compare the standard training method with state-of-the-art deep learning techniques in the context of hybrid modeling. Particularly, the stochastic adaptive moment estimation (ADAM) is considered a landmark in deep learning and was implemented here to train hybrid models. The ADAM method estimates the network parameters, *ω* = {*w,b*}, through the first and second moments of the gradients of the loss function and a set of hyperparameters *α, β*_1_ and *β*_2_, representing the step size and exponential decays of the moment estimations (for details see (Kingma, 2014)). The loss function is the same as in the previous method (Eq. 3). This results in the following implementation:

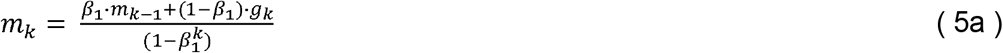

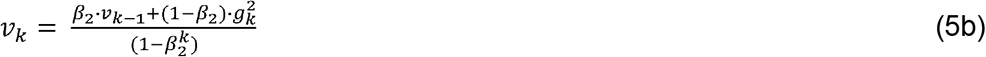

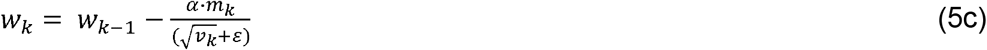

with *k* the iteration number, *m*_*k*_ the first order moment of gradients, *g*_*k*_ the loss function gradients, *v*_*k*_the second order moment of gradients. For the present study, the suggested default parameters of *α* = 0.001, *β*_1_ = 0.9, *β*_2_ = 0.999 and *ε*= 10^−8^ were adopted (Kingma, 2014).

The gradients at each iteration are obtained by solving the sensitivity Eqs. (4a-c). Because the CPU scales exponentially with the size of the network, a different approach to calculate the gradients was explored. Instead of computing the sensitivities of state variables in relation to network parameters, 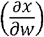, a semidirect approach was implemented where the sensitivities of state variables in relation to network outputs, 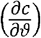, are computed. The semidirect sensitivity equations are as follows (again assuming *y= x*):

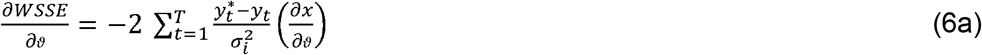

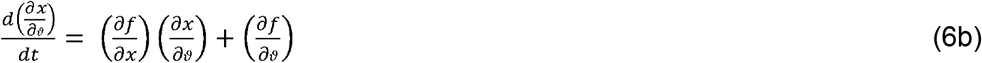

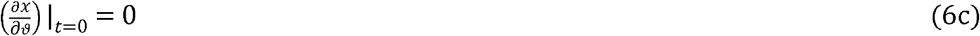

Finally, the loss function gradients 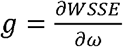 can be computed from the 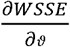 sensitivity (Eq. 6a) by the well-known error backpropagation algorithm through the network (Werbos, 1974). The main advantage of the semidirect method is that the number of ODEs for calculating the sensitivities is massively reduced and independent of the size of the network. This results in a sizable CPU reduction as shown in the results section.

### Case studies

#### *Lee & Ramirez* synthetic data set

A synthetic data set was generated based on the Lee & Ramirez bioreactor model (Lee and Ramirez, 1994). This model is frequently adopted as a benchmark to test different optimal control methods (e.g. (Banga et al., 2005)). The strategy followed in this case study was to train the hybrid models on a information rich data set (time series data) generated by statistical design of experiments, and then to assess the predictive power of the final model in describing the dynamic profiles of the optimal fed-batch obtained by dynamic optimization – optimal control (Lee and Ramirez, 1994).

The Lee&Ramirez model describes the dynamics of biomass (X), substrate concentration (S), inducer concentration (IND), product concentration (P), shock factor (Sh), recovery factor (Re) and reactor volume (V) in a recombinant *Escherichia coli* fed-batch process. Experiments were simulated dynamically for different conditions (see below) applying a Runge-Kutta 4/5^th^ order ODEs solver. Samples were simulated with 1 hour sampling time. Gaussian noise was added to “sampled” variables with standard deviations of 1.5 (X), 5(S) and 0.3(P) (10% of maximum concentration). As shown later in the results section, modeling errors were calculated based on the noisy data (noisy WMSE) and also on the noise free data (noise free WSSE).

A central composite design (CCD) was applied to the process degrees of freedom, namely the induction time between 5-9 hours, pre-induction substrate feed rate between 0-0.8h^-1^, post-induction substrate feed rate between 0-0.8h^-1^ and Inducer feed rate between 0-1h^-1^. This resulted in 25 fed-batch experiments. The 25 fed-batch experiments were included in the training data partition (297 training data points). The validation data set (used only as training stop criterium) was obtained by adding gaussian noise with standard deviations of 1.5 (X), 5(S) and 0.3(P) to the training data set resulting in 297 validation data points. In our experience (results not shown), this partition method maximizes data usage for training and also effectively prevents model overfitting. For the test data set (used to assess the model generalization capacity) the optimal fed-batch with optimized feeding and maximum product concentration of 3.16 g/L (Lee and Ramirez, 1994) was adopted (15 data points). In summary, the models were trained/validated with the 25 DoE experiments and then set to predict the dynamic profiles of the optimal production fed-batch. The optimal production fed-batch delivers a final product mass, which is 34.4% higher than the best DoE fed-batch. The details of the dataset are provided as supplementary material A.

The hybrid model structure adopted for this problem is shown in Figure 2. The reactor has 7 internal sate variables ***x***=[*x, S, P, IND, Sh, Re*]^*T*^ of which only 3 are measured thus ***y***=[*X, S, P*]^T^. The system of ODEs are derived from mass conservation laws and are the same as in (Lee and Ramirez, 1994). The neural network computes 4 reaction terms *ϑ*= [*μ, v*_*S*_, *v*_*P*_, *k*_1_]^*T*^, taken as unknown cellular features that need to be learned from data. The neural network has only 3 inputs ***H***^**0**^ = [*S, IND, Sh*,]^*T*^ which were pre-selected based on prior knowledge of the reaction kinetics for this problem (Lee and Ramirez, 1994).

**Figure 2.**
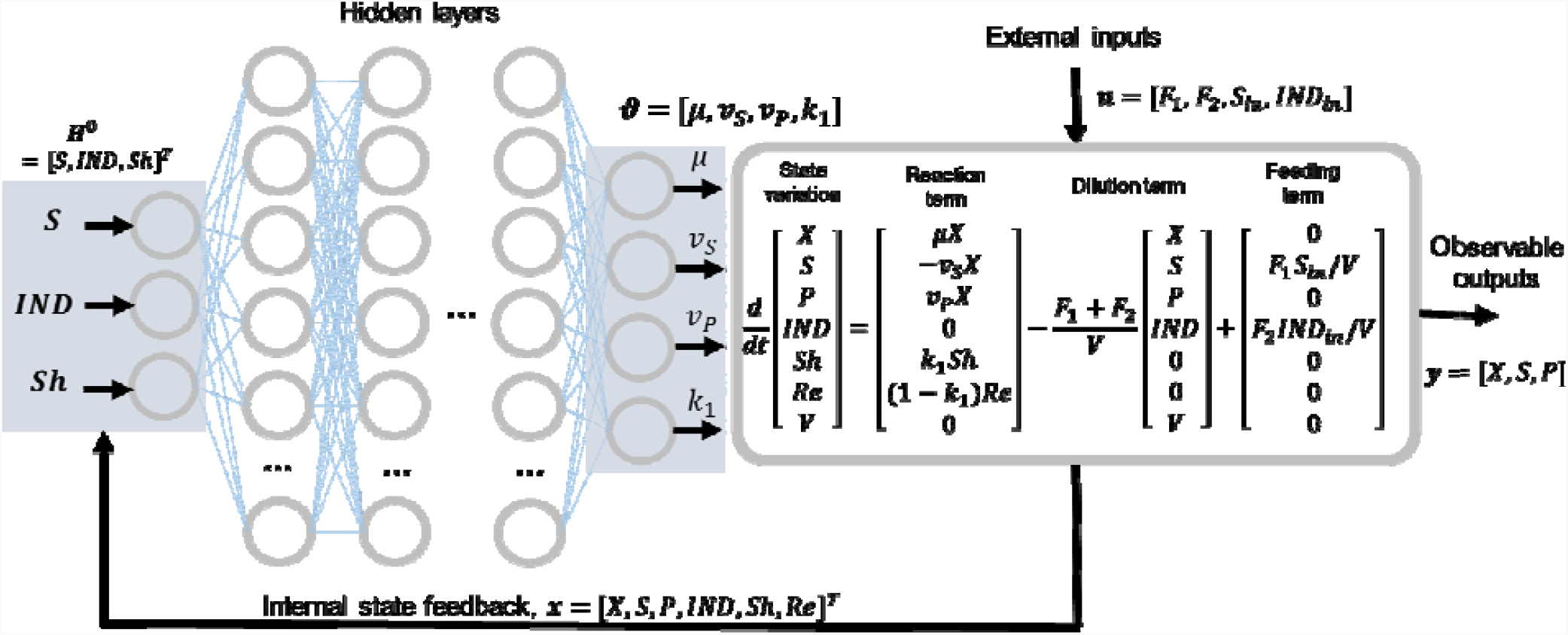
Deep hybrid model structure for the Lee & Ramirez data set. The parametric component is established by a system of ODEs as described in Lee&Ramirez (1994). The specific biologic kinetics are considered mechanistically unknown thus modelled by a deep feedforward network. The job of this model is thus to “learn” from data the biologic kinetics under the constraint of dynamic material balance equations.

#### MUT+ *Pichia pastoris* pilot data set

A MUT+ *Pichia pastoris* expressing a single chain antibody (scFv) was cultivated in a Lab Pilot Fermenter Type LP351, 50 L, Bioengineering, Switzerland with standard instrumentation to measure on-line pH, temperature, pressure, stirrer, airflow and pO2. The wet cell weight and scFv titer were measured off-line. All the details of the experimental procedure are given elsewhere (Teixeira et al., 2006). The reactor operation is divided in three phases: glycerol batch (GB) phase, glycerol fed-batch (GFB) phase and methanol fed-batch (MFB) phase (or post-induction phase). In the GB phase, the initial glycerol level was set at 4%, taking approximately 30 h for complete depletion. Thereupon, the GFB phase starts, following an exponential feeding profile. At the end of the GFB, a transition to the MFB phase is implemented in order to minimize the adaptation time of cells to methanol. After the transition phase, the methanol feeding rate, the pH and the temperature were designed in order to generate process data to optimize scFv productivity (see (Teixeira et al., 2006) for details). A total of 9 experiments were performed with varying methanol feed rate, temperature and pH. In this study, only the MFB phase was considered for hybrid modeling. The data set with the 9 experiments has 207 measurements of biomass wet cell weight in triplicate and 207 measurements of scFv in triplicate. The hybrid model structure adopted for this problem is similar to that of Figure 2 with a few adaptations (discussed later in the results sections).

## RESULTS AND DISCUSSION

### Development of a shallow hybrid model: *Lee & Ramirez* case study

We have first developed a traditional shallow hybrid model for the *Lee & Ramirez* data set. A shallow feedforward network with a single hidden layer with *tanh* activation function was employed. The hybrid model was trained with the standard non-deep method (LMM optimization + cross-validation + random weights initialization from the uniform distribution). The training and validation partition comprehended 25 experiments (825 training patterns) designed by statistical DoE (see methods section). The test partition included a single experiment with the highest protein production (optimal batch obtained by dynamic optimization as reported in (Lee and Ramirez, 1994). The test experiment has a final product mass 34.4% higher than the best training/validation experiment. For a given network size, the training was always repeated 10 times with different weights initialization between [-0.1, 0.1] and only the best result was kept (lower validation error). This procedure was repeated for hybrid models with varying number of nodes in the hidden layer keeping the same data partition and maximum number of iterations of 20000 for comparability. The overall results are shown in Table 1. From these results, it is possible to conclude that the optimal number of hidden nodes is 10 corresponding to the lowest corrected Akaike information criterion (AICc) value (111). Of note, the AICc criterion, which is calculated for the training partition only, coincided with the lowest noise free test error (0.54 noise free WSSE; to note that the noise free WSSE is computed on process data uncorrupted by experimental noise, thus a better metric for accessing the predictive power). This supports the AICc as a suitable hybrid model discrimination criterion in the sense that it discriminates the structure with the highest predictive power. Moreover, the noisy test error of the selected model with 10 hidden nodes (noisy WSSE=1.05) is only moderately higher (11,6%) than the corresponding training error (WSSE=0.941).

**Table 1.** Training results of shallow hybrid models for the Lee&Ramirez data set with 25 training batches (Training WSSE), 25 validation batches (Validation WSSE) and a single test batch with the highest possible productivity obtained by dynamic optimization(Test WSSE noisy/noise free are computed with noisy or noise free target concentrations respectively). The AICc is computed for the training data set only. Each row represents a different model with a given number of hidden nodes (between 1-15) in a single hidden layer with tanh activation function. The hybrid models were trained with the standard non-deep method (LMM optimization with 20000 iterations + cross-validation + random weights initialization between [-0.1, 0.1] from the uniform distribution). The training was repeated 10 times with different weights initialization and only the best result is kept for each model.

| Number of hidden nodes | Training WSSE | Validation WSSE | Test WSSE (noisy) | Test WSSE (noise free) | AICc | CPU time | Number of Weights |
| --- | --- | --- | --- | --- | --- | --- | --- |
| 1 | 20.2 | 20.3 | 42.2 | 2.1 | 2490 | 776 | 10 |
| 2 | 2.57 | 2.77 | 7.53 | 8.12 | 810 | 1320 | 17 |
| 3 | 1.16 | 1.31 | 1.08 | 1.39 | 172 | 1780 | 24 |
| 4 | 1.1 | 1.29 | 1.34 | 1.01 | 146 | 1560 | 31 |
| 5 | 2.77 | 3.07 | 6.56 | 5.42 | 922 | 1390 | 38 |
| 6 | 1.78 | 1.94 | 1.22 | 2.11 | 570 | 1730 | 45 |
| 7 | 1.40 | 1.70 | 7.87 | 7.31 | 389 | 1870 | 52 |
| 8 | 1.09 | 1.32 | 1.14 | 0.76 | 200 | 2050 | 59 |
| 9 | 1.01 | 1.21 | 1.04 | 0.68 | 150 | 2250 | 66 |
| 10 | 0.941 | 1.16 | 1.05 | 0.54 | 111 | 2250 | 73 |
| 11 | 0.949 | 1.22 | 1.33 | 0.83 | 134 | 2360 | 80 |
| 12 | 0.914 | 1.11 | 0.86 | 0.75 | 121 | 2290 | 87 |
| 13 | 0.935 | 1.07 | 1.03 | 0.69 | 154 | 2280 | 94 |
| 14 | 0.944 | 1.15 | 1.10 | 0.93 | 183 | 2230 | 101 |
| 15 | 0.899 | 1.11 | 0.937 | 0.62 | 152 | 2670 | 108 |

### Comparing the deep and shallow hybrid modeling approaches

Several hybrid structures with varying neural network depths (2-4 hidden layers) were compared with the shallow network case (1 hidden layer). The same *Lee&Ramirez* data set and data partition were kept as in the previous section. We first focused on the *tanh* activation (in the hidden layers), which has been the standard for nonlinear regression problems with shallow neural networks (Cybenko, 1989). Every model structure was trained with two different methods: the traditional LMM+CV+tanh and ADAM+CV+tanh. The training was always repeated 10 times and only the best solution (lowest validation error) was kept, as before. The number of iterations of the ADAM method was 20000 as for the LMM method. The overall results are shown in Table 2.

**Table 2.**
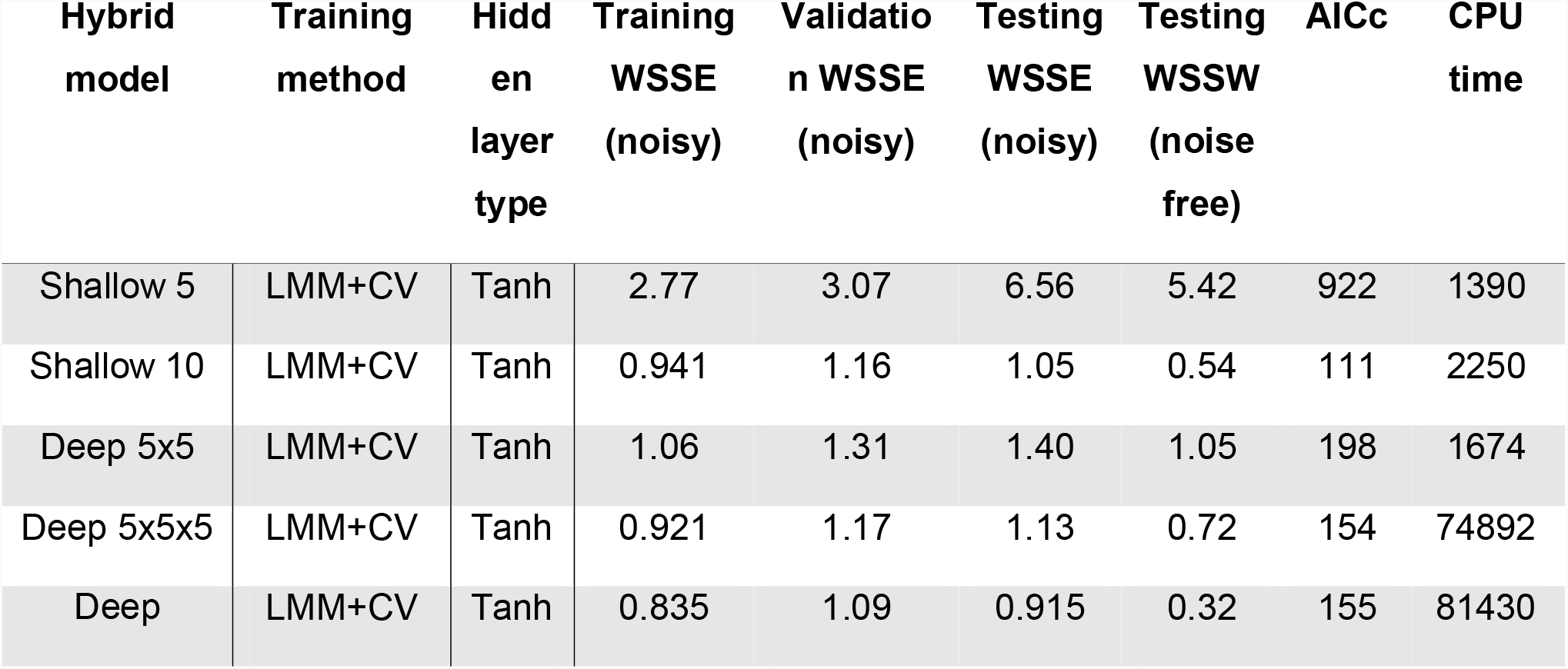
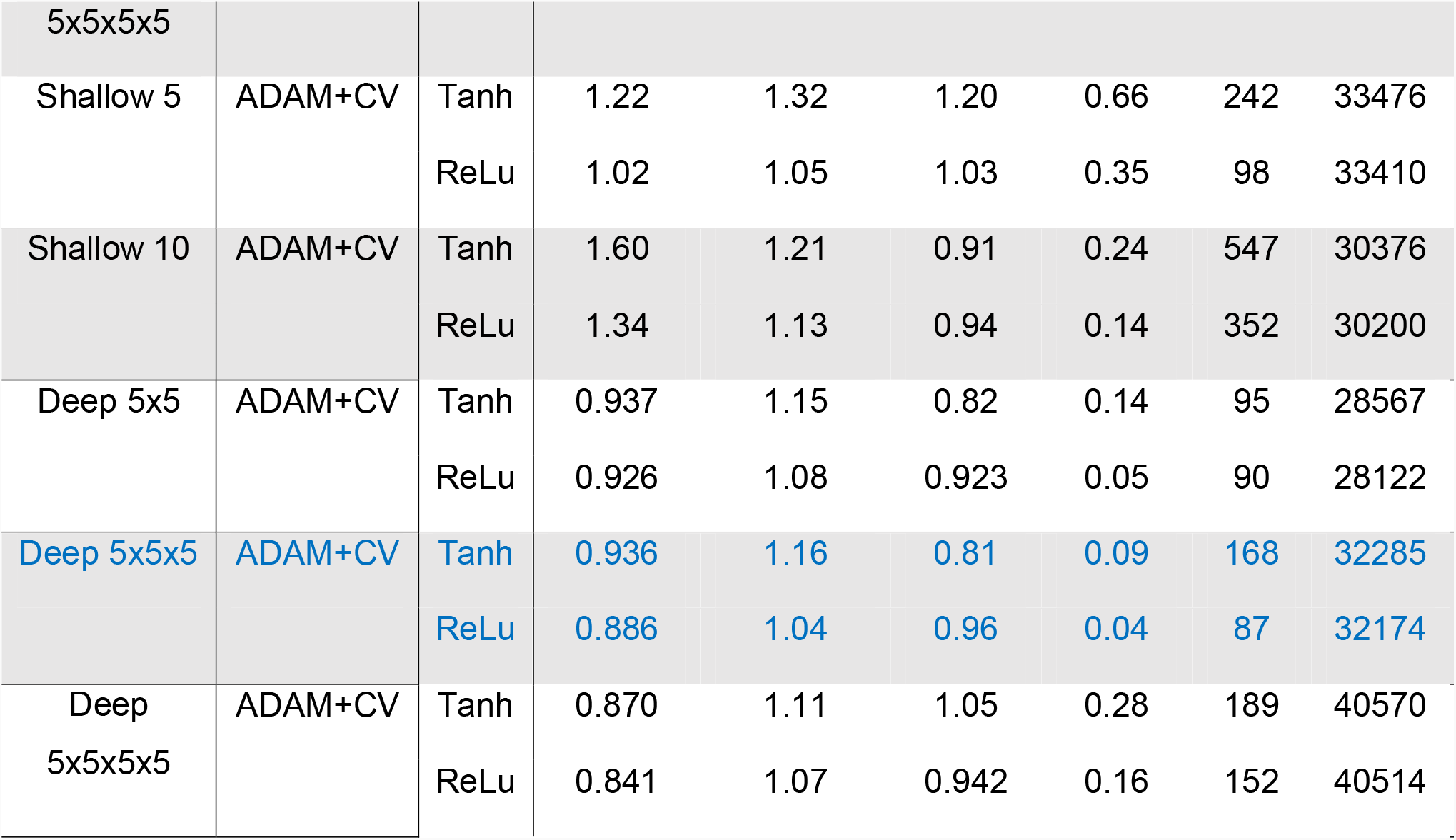
Comparison of deep and shallow hybrid models for the Lee&Ramirez data set (same data partition as in Table 1) trained either by the LMM algorithm or the ADAM algorithm. In all cases cross-validation (CV) and indirect sensitivities were applied. Each row represents a different shallow or deep hybrid model structure using either *tanh* or *ReLU* in the hidden layers. The training was repeated 10 times with different weights initialization and only the best result is kept.

The results in Table 2 clearly show ADAM to outperform the LMM method in what concerns the predictive power of the final model (the noise free test WSSE). This conclusion is valid for deep or shallow hybrid structures. The best shallow structure with 10 hidden nodes (identified in the previous section) improved the noise free test error from 0.54 to 0.24 (>2 fold decrease) with ADAM+CV+tanh. The same conclusions can be taken for the deep structures, without exception. The key conclusion is that the ADAM method systematically increases the predictive power of the final hybrid model for the *Lee&Ramirez* data set.

The best model (with *tanh* activation function) among the deep and shallow structures is the 5×5×5 deep hybrid model with 98 weights, showing a noise free test error (WSSE = 0.09) 2.7 fold lower than the best hybrid shallow case (WSSE=0,24). The AICc miss spotted the discrimination of the best deep model since it identified the 2^nd^ best model (5×5 structure) with, however, comparable performance. In terms of CPU, the ADAM method is generically more expensive than the LMM method for small size networks. This pattern reverses for large size networks (e.g. the best 5×5×5 structure decreased CPU by 2,3 fold with ADAM in comparison to LMM). Thus, the CPU scales more steeply with the network size in the case of LMM training when compared to ADAM training. This generically favors the ADAM method for deep hybrid structures, both in terms of predictive power and required CPU time for training.

Figure 3 shows the effect of the weights initialization on the final training, validation and testing error for the best deep configuration 5×5×5 when the model is trained with LMM or with ADAM. The initial weights values were kept the same for LMM and ADAM training for comparability. Interestingly, the dispersion of the errors for 10 repetitions with different weights initialization is significantly lower for ADAM in comparison to the LMM, irrespective of the data partition (train, validation or testing). Similar results were obtained for the other model configurations (results not shown). There is an outlying point with significantly higher errors for both the LMM and ADAM training. This suggests the ADAM method to be less sensitive to weights initialization. Similar conclusions were reported by (Hiscock, 2019) for standalone deep neural networks, who showed that gradient descent training methods with variable learning rate (such as the ADAM method) are less prone to be trapped in local optima thus less sensitive to weights initialization. The key conclusion to be taken, is that the number of repetitions for different weights initialization may be mitigated in the case of ADAM training. This represents a potential 10 fold cut in CPU time in comparison to the LMM method for the case of 10 repetitions.

**Figure 3.**
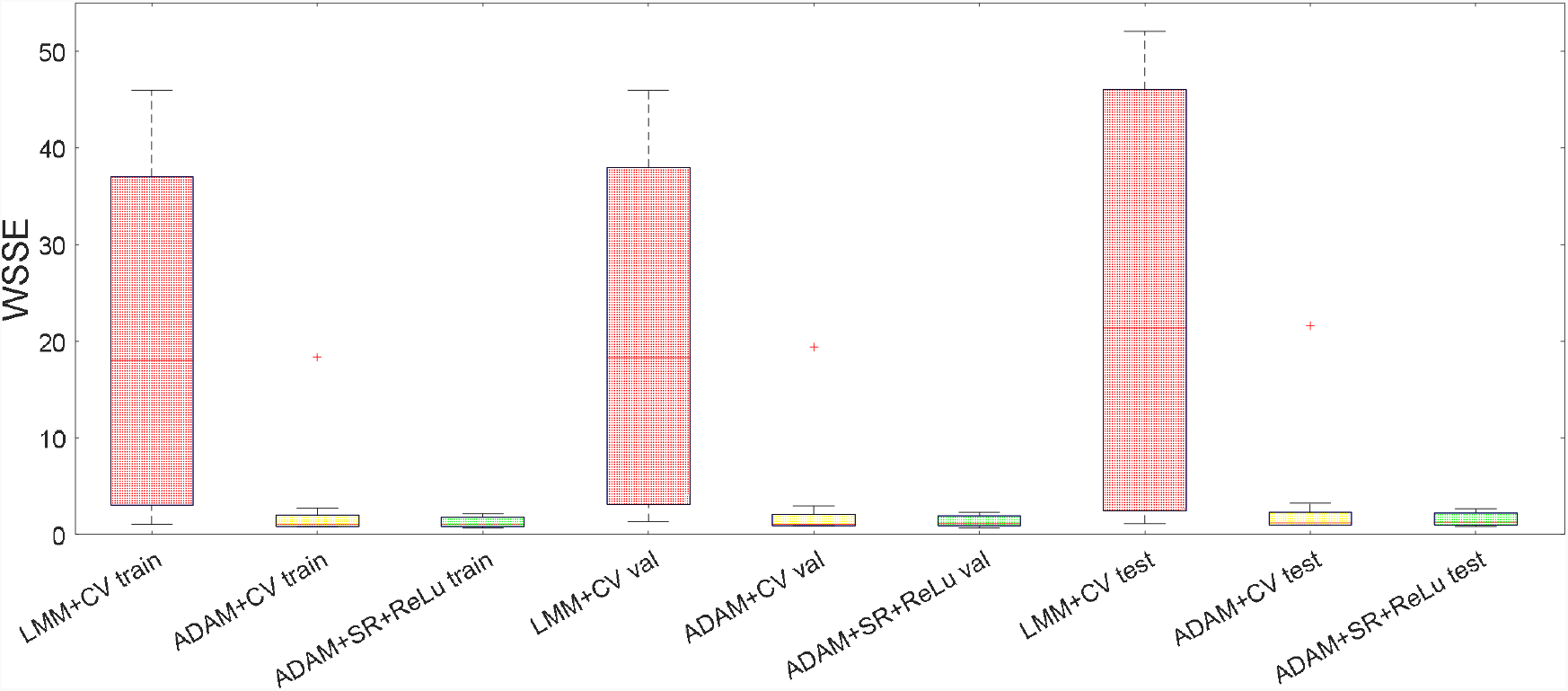
Boxplot of training, validation and testing WSSE for 10 training repetitions of the hybrid deep structure 5×5×5 trained by different training approaches either using the LMM or the ADAM method. Ten sets of initial weights were randomly generated (one per repletion) and kept the same in all tests performed for comparability.

We have further compared two different activation functions in the hidden layers: the *ReLU* and the *tanh* activation functions The *ReLU* function has been a key achievement in deep learning, outperforming the *tanh* function for standalone deep neural networks (Nair and Hinton, 2010). Table 2 compares hybrid model performances using the one or the other activation function in the hidden layers trained by ADAM + CV using the same training procedure. The key conclusion to be taken is that the *ReLU* further improved the training and test error in all cases without exception. The best 5×5×5 structure further decreased the noise free test WSSE from 0.09 (with *tanh*) to 0.04 (with *ReLU*) at comparable CPU cost. Our results clearly show the *ReLU* to be advantageous in a deep hybrid modeling context as previously shown for (standalone) deep neural networks (Nair and Hinton, 2010). The *ReLU* activation function was thus adopted in all proceeding studies.

### Introducing stochastic regularization

Stochastic regularization (SR) has been reported as an effective method to avoid overfitting in deep learning (Srivastava et al., 2014). Here we study the ADAM method with stochastic regularization in replacement of the cross-validation technique. More specifically, ADAM was implemented with the minibatch technique and the weights dropout technique. The minibatch technique consists in a random selection of the training patterns from the uniform distribution using a cutoff probability parameter. Similarly, the weights dropout technique used random weights selection according to a cutoff probability parameter. Figure 4 shows the effect of the minibatch size probability and of the weights dropout probability on the testing error for the deep configuration 5×5×5. The hybrid 5×5×5 model was trained 10 times with different weights initialization with varying minibatch and weights dropout probabilities. Figure 4 shows the lowest WSSE test among the 10 repetitions as function of the minibatch size probability and of the weights dropout probability. The training performance is indeed very sensitive to the choice of these two parameters. The optimal minibatch probability is ∼90% and the optimal dropout probability is ∼50%. The final noise free test WSSE was 0.0258, which is 35.5% lower than the corresponding solution without stochastic regularization (Table 2, ADAM+CV+ReLU). The final train and test errors among the 10 repetitions are shown in Figure 3. Interestingly the stochastic regularization eliminated the outlying training result obtained by LMM+CV and ADAM+CV in the previous section. This result is promising because it shows the weights initialization to have practically no influence on the final training outcome. If repetitions are not needed, the CPU cost may be significantly reduced in relation to the LMM+CV or ADAM+CV methods.

**Figure 4.**
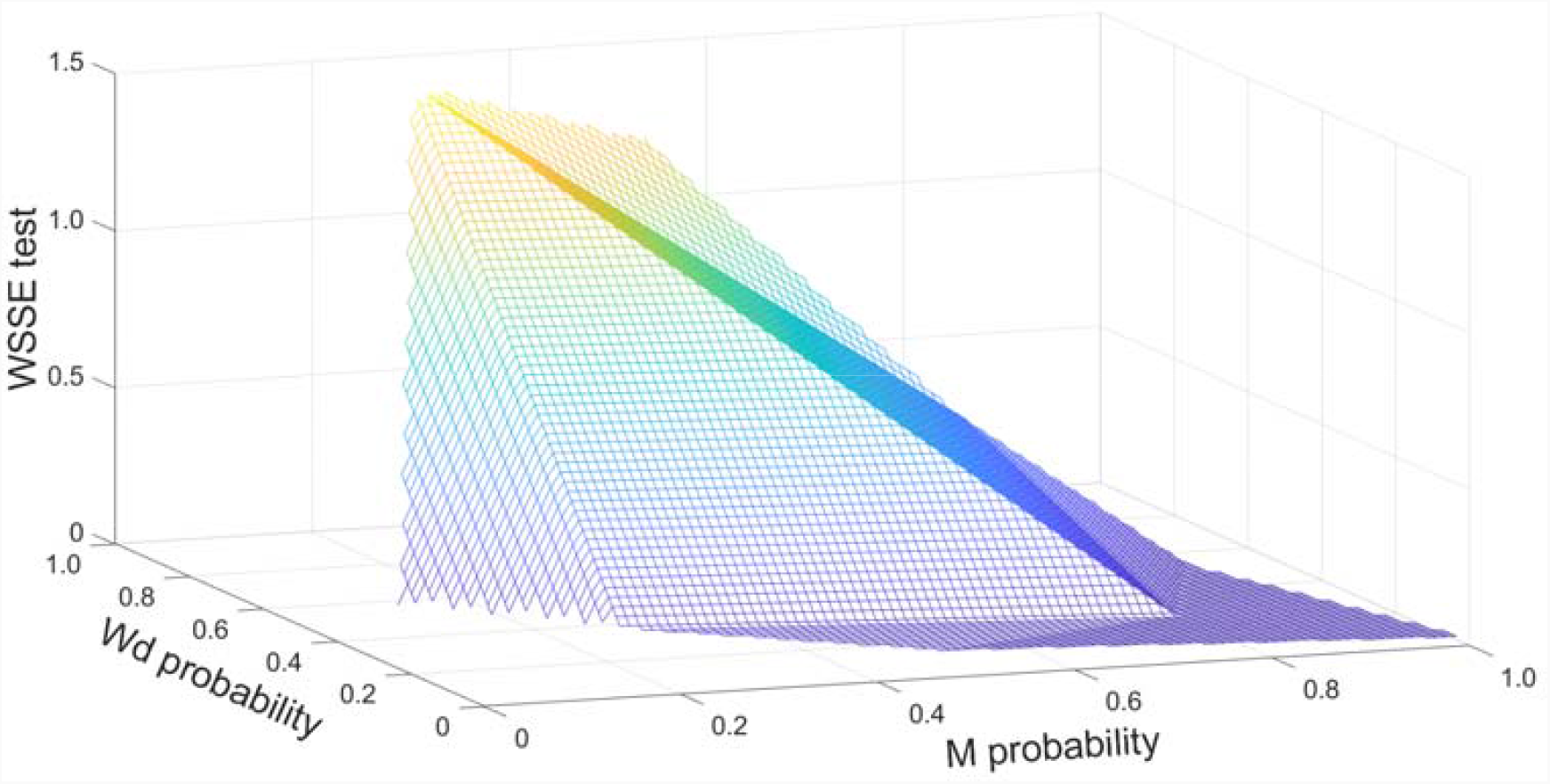
Effect of stochastic regularization (SR) on the predictive power of the hybrid model configuration 5×5×5 trained with ADAM + SR + indirect sensitivities with 20000 iterations for the Lee&Ramirez data set. Obtained noise free test WSSE over minibatch probability (M probability) and weights dropout probability (Wd probability).

### Speeding up hybrid deep learning by semidirect sensitivities

The results above support ADAM + deep networks + stochastic regularization to produced hybrid models with higher predictive power in comparison to the traditional shallow hybrid approach. Nevertheless, deep models tend to have large networks with the CPU time increasing with the network size (Luo et al., 2005). Solving the sensitivity equations is responsible for a significant part of the CPU cost. Taking the 5×5×5 hybrid structure as example, solving the sensitivity equations implies integrating 98×5 = 490 ODEs along with the hybrid model ODEs for the computation of the objective function and objective function gradients. Such a large number of ODEs represents a significant CPU burden. We have investigated a different implementation of the sensitivity method, namely the semidirect sensitivity equations (see methods section) in an attempt to reduce CPU time. In the semidirect approach, a much lower number of 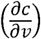 sensitivity equations are integrated over time. For the same 5×5×5 hybrid structure, the 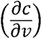 sensitivities only require 5×4=20 ODEs to be integrated over time. Furthermore, the semidirect sensitivity equations are independent of the network size (they depend only on the number of states and on the number of network outputs). Figure 5 shows the variation of the train and test cost function over CPU for the configuration 5×5×5. This result shows that the semidirect sensitivity equations produced a comparable final training WSSE in relation to the indirect sensitivity equations. The convergence is however much faster. The CPU time could be reduced by 77.4% when adopting the semidirect sensitivity equations in comparison with the indirect approach. Furthermore, the test error follows similar patterns for both methods reaching a comparable final value.

**Figure 5.**
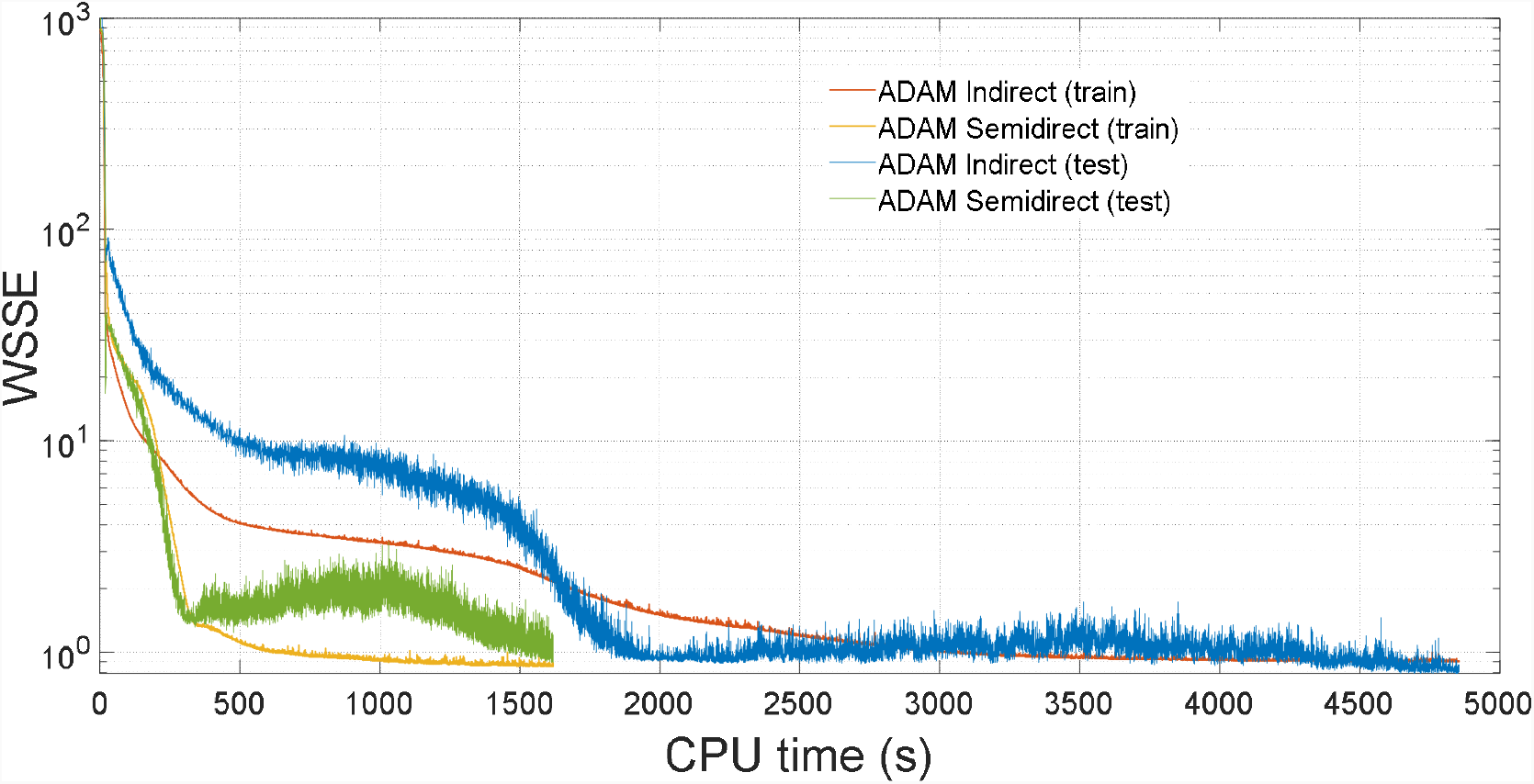
Training and testing error (WSSE) over CPU time for 1) shallow hybrid model {10} + LMM +CV with ten repetitions (blue line) 2) the hybrid model 5×5×5 trained with ADAM + stochastic regularization + indirect sensitivities (red line) and 3) ADAM + stochastic regularization + semidirect sensitivities (yellow line)

Figure 6 shows the prediction of the optimal batch dynamics by the hybrid 5×5×5 model trained with ADAM+SR+ReLU+semidirect compared to the standard shallow model with 10 hidden nodes (LMM+CV+tanh+indirect). The noise free test WSSE was 0.03 and 0.54, respectively (94,4% reduction). It may be seen that both modes are able to describe fairly well the dynamics of the test experiment up to 7.5 hours. There are however some visible differences towards the end of the cultivation. The shallow hybrid model underestimated the final biomass and final product by 15.3% and 13.8% respectively, whereas the deep hybrid model overestimated the final biomass by 2.7% and underestimated the final product by 5.8%.

**Figure 6.**
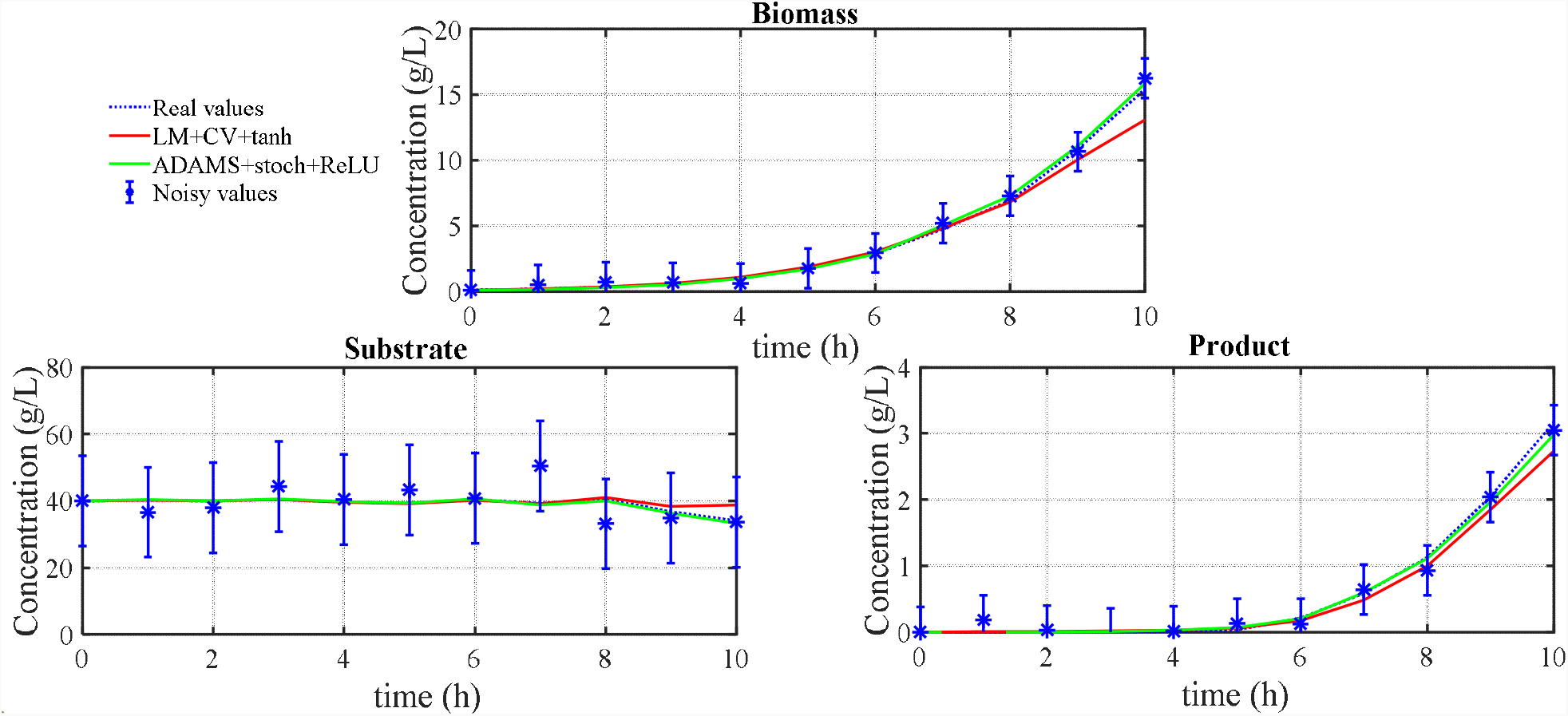
Prediction of the dynamic profiles of observable variables (biomass -X, substrate– S and product–P) by the shallow (10) hybrid model and by the deep (5×5×5) hybrid model for the test batch of the Lee&Ramirez data set. Circles represented observations and respective ± standard deviation. The dashed line represents the “true” process behavior (hidden to the training of the hybrid models). The blue line represents the predictions of the shallow hybrid model. The red line represents the prediction by the deep hybrid model. The shallow hybrid model used the tanh function and was trained by the traditional non-deep method (LMM algorithm + CV + indirect sensitivities + 10 repetitions and only the best result is kept). The deep hybrid model used the ReLU activation function and was trained by the novel method (ADAM + SR + semidirect sensitivities + no repetitions).

### Pilot scale *Pichia pastoris* case study

The two best shallow and deep hybrid structures previously identified for the Lee&Ramirez case study (namely the shallow structure with 10 nodes in the hidden layer and the deep 5×5×5 structure) were applied for the *Pichia pastoris* case study. The biomass and product material balance equations, and the shock factor and recovery factor ODEs are kept the same in both models. A few modifications were however required as follows:

⍰ The inducer material balance equation was removed because in the MUT+ *P. pastoris* expression system the methanol is simultaneously the main carbon source and the inducer of foreign protein expression.
⍰ The substrate material balance equation was also removed because methanol concentration (the substrate) was not measured. This is a limitation imposed by the experimental protocol. Instead, the measured volumetric methanol feed rate (*F*_*met*_, g/Lh) and the measured total methanol fed to the reactor (g) were set as external inputs to the neural network.
⍰ Temperature (T) and pH were also added as external inputs to the neural network as these two parameters varied between 17.2-30.1ºC and pH 4.0-7.0 in the experiments performed as part of a design of experiments to study the influence of these two parameters in the protein expression.
⍰ The neural network computed the volumetric protein production rate (output) instead of the specific protein production rate as in the case of *Lee&Ramirez*. It is known that *Pichia pastoris* secretes proteases that hydrolyses the target product on certain experimental conditions (Cereghino and Cregg, 2000). The neural network is thus set to calculate the apparent volumetric production rate of the scFv which lumps the synthesis and hydrolysis in the same kinetic term.

The number of parameters in both models is the same, namely 123. The shallow hybrid structure was trained with the traditional method (LM+CV+*tanh*+direct, 10 repetitions with random weights initialization from the uniform distribution) whereas the deep hybrid structure was trained with the new method (ADAM+SR+ReLU+semidirect, weight dropout probability of 0.5, minibatch probability of 0.9 and no repetitions). Eight reactor experiments were used for training-validation (validation data points were obtained by adding gaussian noise to the training data points as in the Lee&Ramirez case study) and just one experiment for testing. All possible training-validation/testing permutations were evaluated. The overall results are shown in Table 3 where each row represents a different training-validation/testing permutation. As illustrative example, Figure 7 shows the measured and predicted dynamic profiles of biomass and product for the case of experiment F66 used for testing. The key conclusions to be taken is that both the training and testing WSSEs were lower for the deep hybrid structure in relation to the shallow structure, in all data partitions tested without exception.

**Table 3.** Comparison of deep and shallow hybrid models for the pilot reactor MUT+ *Pichia pastoris* data set. Each row represents a hybrid model obtained by training over a different training/testing data permutation (Test batch ID refers to the batch used for testing while the remaining 8 batches were used for training/validation). Shallow hybrid models tanh activation function and were trained by the traditional non-deep method (LMM algorithm + CV + indirect sensitivities + 10 repetitions and only the best result is kept). Deep hybrid models used the reLU activation function and were trained by the novel method (ADAM + SR + semidirect sensitivities + no repetitions).

| Test batch ID | Model type | Training WSSE (noisy) | Testing WSSE (noisy) | AICc | CPU time |
| --- | --- | --- | --- | --- | --- |
| F037 | Shallow 10 | 2.18 | 2.58 | 664 | 19560 |
|  | Deep 5x5x5 | 1.79 | 2.13 | 587 | 13980 |
| F044 | Shallow 10 | 2.42 | 3.94 | 700 | 46440 |
|  | Deep 5x5x5 | 2.14 | 3.73 | 633 | 19980 |
| F048 | Shallow 10 | 2.01 | 2.55 | 626 | 15060 |
|  | Deep 5x5x5 | 1.96 | 2.28 | 618 | 12000 |
| F061 | Shallow 10 | 2.65 | 4.69 | 738 | 22860 |
|  | Deep 5x5x5 | 1.98 | 4.05 | 620 | 14520 |
| F066 | Shallow 10 | 2.54 | 2.82 | 722 | 13680 |
|  | Deep 5x5x5 | 1.59 | 1.86 | 542 | 9660 |
| F007 | Shallow 10 | 2.79 | 4.13 | 752 | 23640 |
|  | Deep 5x5x5 | 2.24 | 2.98 | 663 | 13320 |
| F009 | Shallow 10 | 2.82 | 4.72 | 754 | 30180 |
|  | Deep 5x5x5 | 2.62 | 3.76 | 730 | 12480 |
| F018 | Shallow 10 | 2.48 | 3.28 | 710 | 15900 |
|  | Deep 5x5x5 | 2.31 | 2.67 | 684 | 10200 |
| F072 | Shallow 10 | 3.15 | 4.85 | 791 | 26100 |
|  | Deep 5x5x5 | 3.01 | 3.98 | 775 | 14820 |
| **Sum** | **Shallow 10** | **23.04** | **33.6** | **6457** | **213420** |
|  | **Deep 5x5x5** | **19.64** | **27.4** | **5852** | **120960** |

**Figure 7:**
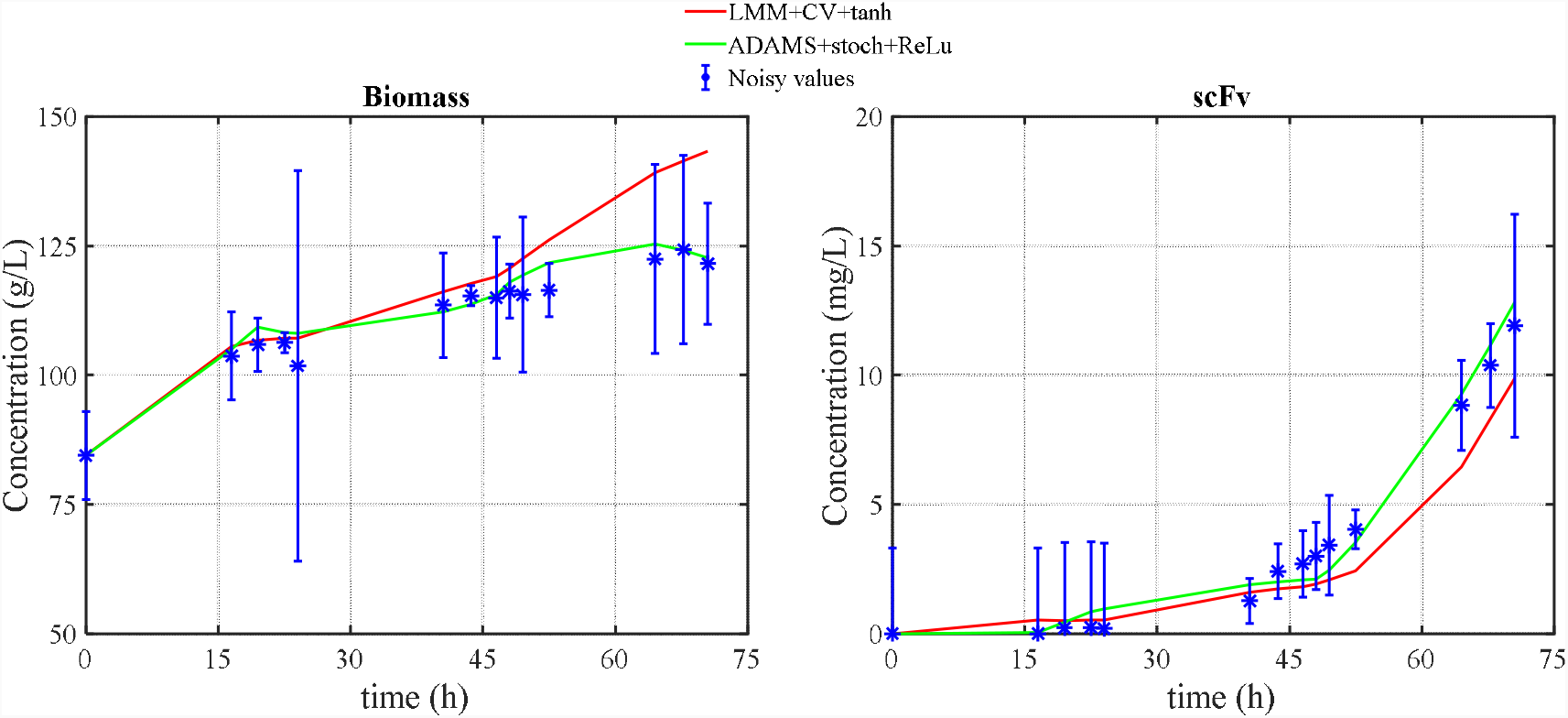
Prediction of the dynamic profiles of observable variables (biomass-X, and product–scFv) by the shallow (10) hybrid model and by the deep (5×5×5) hybrid model for the test batch F066 of the MUT+ *Pichia pastoris* pilot data set. Circles represent observations and respective ± standard deviation. The blue line represents the predictions of the shallow hybrid model. The red line represents the prediction of the deep hybrid model. The shallow hybrid model used the tanh activation function and was trained by the traditional non-deep method (LMM algorithm + CV + indirect sensitivities + 10 repetitions and only the best result is kept). The deep hybrid model used the ReLU activation function and was trained by the novel method (ADAM + SR + semidirect sensitivities + no repetitions).

The differences between the dynamic profiles of biomass and scFv are clearly visible in Figure 7. The predicted final scFv titer by the shallow hybrid model is 17.5% below the experimental value whereas the deep hybrid model overestimated the experimental value by 4.2% only. Taking all data partitions together (last row in Table 3), the average training WSSE decreased by 14.8% whereas the average testing WSSE decreased by 18.4% for the deep hybrid structure in relation to the shallow hybrid structure. Moreover, the average CPU time decrease by 43.4% when applying the deep methodology in comparison to the standard methodology.

## CONCLUSIONS

In this study the general bioreactor hybrid model was revisited and some of the most recent deep learning techniques were investigated in the context of hybrid modeling, where neural networks are combined with systems of ODEs. The effect of increasing the depth of the neural network resorting to two different training approaches was investigated. The traditional approach uses the Levenberg-Marquardt optimization coupled with the indirect sensitivities, cross-validation and *tanh* activation function. The novel hybrid deep approach uses the adaptive moment estimation method (ADAM), semidirect sensitivities, stochastic regularization and ReLU activation functions in the hidden layers. Two applications were addressed, one with a synthetic data set, the other with an experimental dataset collected in a pilot 50 L bioreactor. The key conclusion to be taken is that there is a clear advantage of adopting hybrid deep models both in terms of predictive power and in terms of computational cost in relation to the shallow hybrid case. In the *Lee&Ramirez* case study, the prediction error decreased 94.4% and the CPU decreased 29%. In the case of the *P. Pastoris* case study, the prediction error decreased 18.4% and the CPU decreased 43,3%. The ADAM method coupled with stochastic regularization shows two significant advantages. First, it is practically insensitive to weight initialization thereby eliminating the need for training repetitions. Second, the stochastic nature of the method is less sensitive to experimental noise eliminating the need for cross-validation. Lastly, the introduction of semidirect sensitives, further decreases the CPU time particularly for large deep structures as the number of sensitivity equations (that need to be integrated over time) becomes independent of the number of hidden layers.

## Acknowledgments

JP acknowledges PhD grant SFRD/BD14610472019, FCT, Fundação para a Ciência e Tecnologia, Portugal and RSC the contract CEECIND/01399/2017. This work has received funding from the European Union’s Horizon 2020 research and innovation program under grant agreement No 101000733 (POMICON).

